# Genetic elements orchestrating *Lactobacillus crispatus* glycogen metabolism in the vagina

**DOI:** 10.1101/2022.04.23.489263

**Authors:** Rosanne Hertzberger, Ali May, Gertjan Kramer, Isabelle van Vondelen, Douwe Molenaar, Remco Kort

## Abstract

Glycogen in the female lower reproductive tract is a major carbon source for vaginal colonization and acidification by common vaginal *Lactobacillus* species, such as *Lactobacillus crispatus*. Previously we identified the pullulanase gene *pulA in Lactobacillus crispatus*, correlating with its ability to autonomously utilize glycogen for growth. Here we further characterize genetic variation and differential regulation of *pulA* affecting the presence of its gene product on the outer surface layer. We show that alpha-glucan degrading activity dissipates when *Lactobacilllus crispatus* is grown on glucose, maltose and maltotriose, in agreement with carbon catabolite repression elements flanking the *pulA* gene. Proteome analysis of the S-layer confirmed that the pullulanase protein is highly abundant in an S-layer enriched fraction, but not in a strain with a defective pullulanase variant or in a pullulanase-sufficient strain grown on glucose. In addition, we provide evidence that *Lactobacillus crispatus pulA* mutants are relevant *in vivo*, as they are commonly observed in metagenome datasets of human vaginal microbial communities. Analysis of the largest publicly available human vaginal metagenome dataset indicates that 15 out of 272 samples, containing a *Lactobacillus crispatus pulA* gene, contain a defective variant of this gene. Another 23 out of 272 samples show large deletions or transposon insertions. Taken together, these results demonstrate that both environmental as well as genetic factors explain variation of *Lactobacillus crispatus* alpha-glucosidases in the vaginal environment.

## 1. Introduction

The vaginal environment of reproductive-age women is unique amongst vertebrates in several aspects: it has a low pH due to high levels of lactate (∼100-150 mM) [1] it has abundant glycogen levels [2] and its microbial community is dominated by *Lactobacillus* species [3,4]. In recent years more evidence has emerged that the vaginal microbiome is associated with reproductive and sexual health. Women with a *Lactobacillus crispatus* microbiome are at lower risk of vaginal infections and sexually communicable infections [5– 7]. In addition, vaginal *Lactobacillus crispatus* is inversely associated with more serious health outcomes such as subfertility [8,9], preterm birth [10] and persistent HPV infection [11,12].

We previously demonstrated that most vaginal *L. crispatus* isolates are capable of autonomous growth on glycogen. A disrupted N-terminal signal peptide sequence in the pullulanase gene, either by small mutations or structural variants, coincided with the inability of growing on glycogen [13].

A recent key study confirmed alpha-glucosidase activity of pullulanase genes from different vaginal bacteria by heterologous expression. It showed that the *Lactobacillus crispatus* pullulanase degrades glycogen, pullulan and amylose retaining activity at pH values that are common in a *Lactobacillus*-dominated vaginal environment [14]. *L. crispatus* strains isolated from low-glycogen environments, such as the human or poultry gut, more often lacked the pullulanase gene indicating niche-specificity [15].

An open question in the field is the role of vaginal glycogen stores in bacterial acidification and the origin of vaginal glycogen degrading enzymes. Vaginal samples show alpha-glucosidase activity, which was postulated to be mostly of human origin on the basis of antibody [16,17] and proteomic analysis [18]. At the same time results from two studies have indicated that alpha-glucosidases in a subset of vaginal samples may be of bacterial origin on the basis of optimal pH and substrate and product profiling [14,19]. However, a metaproteomic study of vaginal lavages from women with a *crispatus*-dominated microbiome, failed to detect any *L. crispatus* pullulanase protein, which poses questions about host-related or genetic factors influencing its presence [18].

Here we present results showing substantial disruption and down-regulation of the *Lactobacillus crispatus* pullulanase gene. We use a mass spectrometric approach to demonstrate the presence of the *Lactobacillus crispatus* pullulanase enzyme in an S-layer enriched fraction, its downregulation in glucose-, maltose- and maltotriose-grown cultures and its absence in a strain with a defective *pulA* gene. Analysis of a vaginal metagenomic dataset demonstrates the presence of a significant number of mutations and structural variants expected to fully disrupt pullulanase functionality *in vivo*, indicating that both environmental as well as genetic factors may refrain vaginal *Lactobacillus* from metabolizing glycogen.

## 2. Results

### 2.1 Substrate utilization of pullulanase sufficient and deficient Lactobacillus crispatus strains

Previously we reported a collection of *Lactobacillus crispatus* strains with variable growth on glycogen associated with the presence of an intact amino-terminal signal peptide for protein secretion in the pullulanase gene product (PulA) [13]. Here we further explore the substrate range of a *Lactobacillus crispatus* deficient (RL09) and sufficient (RL10) strain.

To verify the identified deletion of two nucleotides we used a PCR-amplified 300 bp DNA fragment encoding the PulA N-terminal region (including 42 nucleotides upstream the start site) of strains RL09 and RL10 and resequenced with Sanger dideoxy sequencing. This confirmed the presence of the deletion of two nucleotides in the signal peptide sequence of RL09 (hereafter referred to as *pulA*^*-*^ compared to RL10 (hereafter referred to as *pulA*^*+*^ (Appendix Figure 1A).

**Figure 1.**
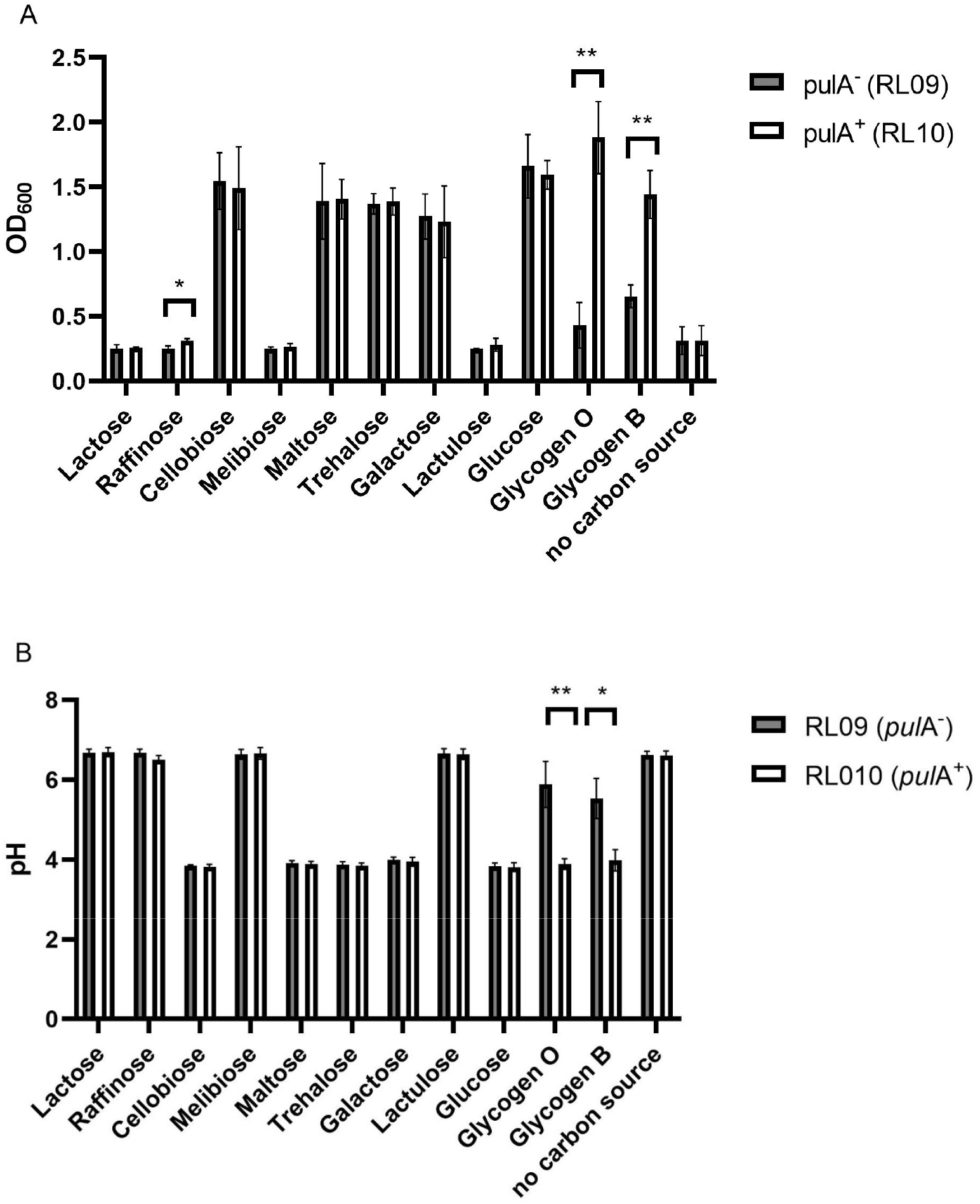
Carbohydrate utilization by *Lactobacillus crispatus* strains *pulA*-(RL09) and *pulA*^*+*^ (RL10). A Optical density (600nm) and B pH after serial propagation in NYCIII medium supplemented with different carbohydrates. The bars represent the mean of 3 biological replicates, the standard deviations are shown as error bars. * p < 0.05, ** p < 0.005 (multiple t-test with false discovery rate).

Previously we found *pulA*^-^ strain RL09 unable to grow on glycogen. We reassessed growth and acidification of this strain and *pulA*^*+*^ strain RL10 on NYCIII medium without glucose supplemented with 11 different carbohydrates to analyze other differences between the strains in metabolic profile. Optical density and medium pH after 24 hours of anaerobic growth are shown in Figures 1A and B, respectively.

Both *L. crispatus* strains were equally capable of growth on all carbon sources except for lactose, raffinose, cellobiose and lactulose, which is in disagreement with previous findings [20] Growth was accompanied by acidification of the medium to a pH of 3.9, corresponding to the *in vivo* vaginal pH [1] and the pKa of lactic acid. Glycogen, both of oyster and bovine origin, was the only carbohydrate resulting in a clear difference between the strains: the pullulanase-deficient strain grew to an optical density of ∼0.5 and acidified to a pH of 6.0 (Oyster glycogen) and 5.5 (Bovine glycogen) possibly due the presence of smaller maltodextrins and glucose after autoclaving the glycogen. The growth of the pullulanase-sufficient strain on glycogen was comparable to that on glucose, acidifying the medium to a pH of 3.9. Regarding growth on the 11 carbohydrates tested, we conclude that *pulA*^*+*^ and *pulA*^*-*^ strains do not substantially differ in their metabolic capacities apart from the ability to grow on glycogen.

### 2.2 Starch degradation activity of pullulanase sufficient and deficient Lactobacillus crispatus strains on various carbon sources

Next we measured enzymatic activity of the pullulanase enzyme to understand its regulation and to find pullulanase-expressing conditions that would allow for a direct proteomic comparison between the two strains. Genetic elements in the locus of the *pulA* gene are strongly indicative of carbon catabolite repression, similar to the regulation of pullulanase in other lactobacilli [21]. The promoter region of the gene has a *cre* (catabolite responsive element) like palindromic sequence (TGTTATCGATAACA). The catabolite responsive element contains a well-known binding site for the global regulator CcpA (carbon catabolite protein A) suppressing metabolic pathways for alternative carbohydrates in the presence of glucose in various *Lactobacillus* species [22–24]. Downstream of the *pulA* gene we identified two open reading frames with homology to the LacI family of repressors, containing a DNA binding domain in one open reading frame (1-110) and an effector domain (91-219) in the other, possibly involved in catabolite repression through binding of breakdown products of glycogen (see below).

To verify the predicted regulation by carbon source we grew the pullulanase-sufficient strain *L. crispatus* to stationary phase on NYCIII medium supplemented with glycogen, maltotriose, maltose, glucose or galactose and measured alpha-glucosidase activity in the pellet and supernatant. We included galactose since lactic acid bacteria generally show preference of glucose over galactose [25,26] and we hypothesize that galactose does not repress *pulA* expression to the same extent as glucose.

After 48 hours of growth, spent medium and bacterial cells were mixed with starch solution to detect starch degradation in a standard starch-iodine assay. Starch was used as a substitute for glycogen, since it can be easily detected using iodine and has an amylopectin moiety with alpha1-4 and alpha1-6 linkages comparable to glycogen. Total starch degradation for a period of 24 hours at 37 degrees was used as a measure of alpha-glucosidase activity in the sample. Starch degradation was observed in pellet (Figure 3) as well as spent growth media (Appendix figure A2) of *L. crispatus pulA*^*+*^ strain RL10 after growth on glycogen and galactose, but not after growth on glucose, maltose and maltotriose, confirming the repression of this gene under these conditions.

Alpha-glucosidase activity in cells and spent medium of the *pulA*^+^ strain was comparable to the *pulA*^-^ strain on all substrates except for glycogen since the *pulA*^-^ strain did not sufficiently grow on glycogen to measure starch degradation. Bacterial cells or supernatants of the *pulA*^-^ strain grown on glucose, maltose, maltotriose or galactose did not degrade any starch conforming the link between the defective pullulanase enzyme and glycogen metabolism.

### 2.3 S-layer enrichment and proteome analysis

In our previous study we identified a surface layer associated protein (SLAP) domain in the *Lactobacillus crispatus* pullulanase protein sequence [13], which prompted us to investigate its localization in the surface layer. We hypothesized that the natural variation in the amino-terminal signal peptide for secretion of pullulanase would result in the presence or absence of an active pullulanase in the S-layer. Furthermore, we wanted to confirm that the catabolite repression of pullulanase by glucose, as observed in this study by use of the starch degradation assay (Figure 2), would be reflected by the levels of pullulanase in the S-layer. We utilized galactose as a carbon source since it 1) allows for growth of both *pulA*^*-*^ and *pulA*^*+*^ strains and 2) is a condition under which we observed cell-associated starch degradation activity (figure 2). We compared cells of the *pulA*^*+*^ RL10 strain grown on galactose, glucose and glycogen in order to verify expression of pullulanase in an S-layer enriched fraction under these conditions.

**Figure 2.**
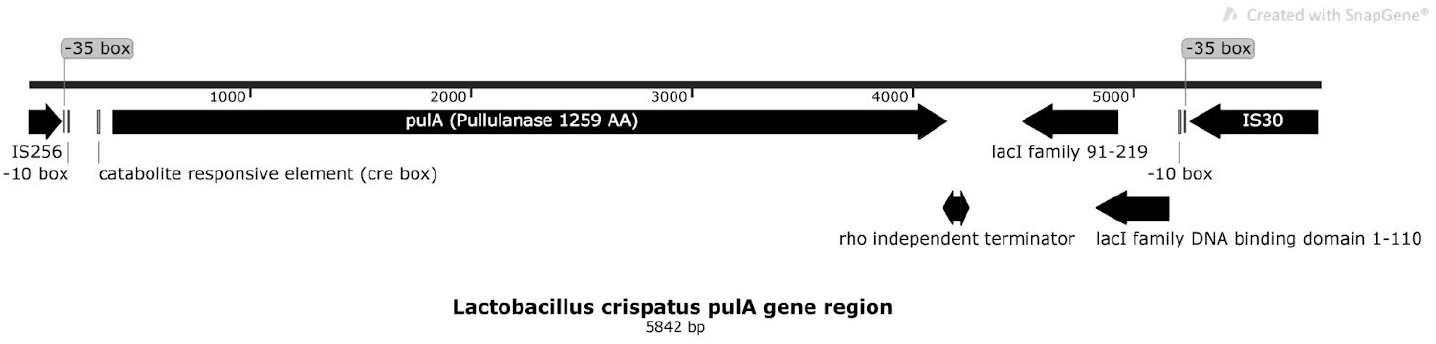
The *pulA* locus (4842 bp) in the chromosome of *Lactobacillus crispatus* RL10.

**Figure 3.**
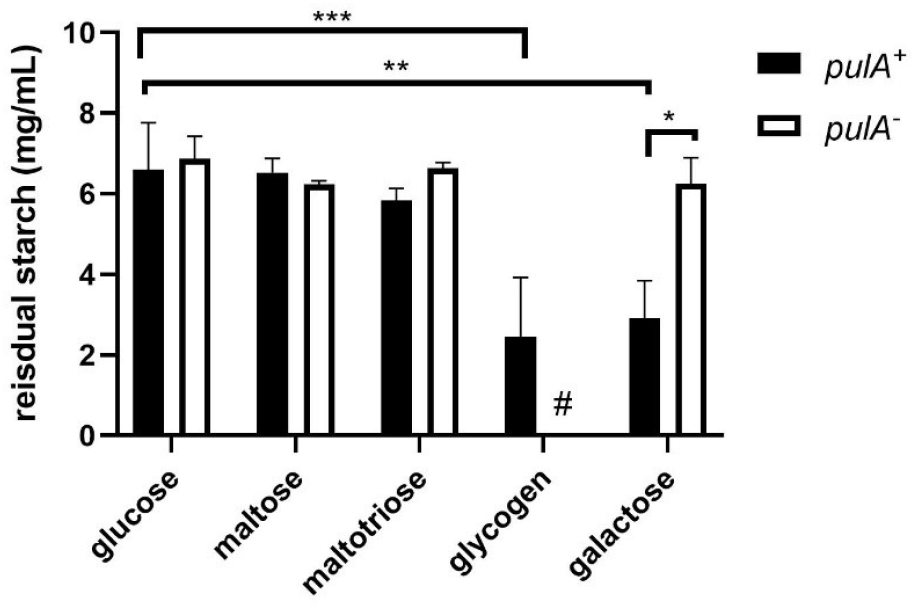
Cell-bound starch degrading activity of *Lactobacillus* crispatus pullulanase sufficient and deficient strains grown on NYC-, glucose, maltose, maltotriose, glycogen (only *pulA*^*+*^) and galactose. Cells were cultured in NYCIII medium supplemented with these carbohydrates and after centrifugation and resuspension of the pellet incubated in a starch solution (7.5 g/L). Asterisks indicate *p*-values calculated by Mann-Whitney of * p<0,05, ** p<0,005, *** p<0,0005. Hashtag indicates no data due to the absence of growth of this strain on glycogen.

The S-layers form the outermost structure of the cell envelope and represent up to 15% of the total protein content [27]. As the S-layer protein subunits (SLPs) and S-layer associated proteins (SLAPs) are non-covalently linked to the cell wall, we released the S-layer fraction from the cells and let it disintegrated into monomeric polypeptides by exposure of the cells to high concentrations of the denaturing agent LiCl. Mass-spectrometric analysis of this isolated S-layer fraction led to the identification of peptides of at least 28 SLAP-domain containing proteins, including a number of abundant SlpA–domain containing proteins (Table 1). The most prominent distinction in the amount of all proteins identified in RL09 and RL10 S-layers is a 90-fold difference for pullulanase (also known as alpha-dextrin endo-1,6-alpha-glucosidase). The minor background of pullulanase peptides in RL09 may result from lysed cells releasing intracellular pullulanase. S-layer fractions of the *pulA*^+^ strain RL10 grown on glucose showed a 24-fold lower pullulanase presence compared to fractions from glycogen grown cells. Two uncharacterized surface layer proteins (A0A7T1TNA6 and A0A125P6L8) are strongly upregulated in the presence of glucose by a factor of over 100; all other proteins show minor differences in levels with ratios between 0.2 and 2.7. The overall fraction of S-layer proteins appears to have been reduced in the presence of glucose (0.2 *versus* 0.4-0.5 in the other three fractions).

**Table 1.** Levels of SLAP domain containing proteins in the S-layer enriched fraction of *Lactobacillus crispatus*. Columns in the table include ACC NO, UniProt Accession Number; PEPT, number of detected peptides; RL09 GAL, expression levels (iBAQ) in *L. crispatus pulA-*strain RL09 cultivated on galactose as only carbon source, RL10 GAL, expression levels in *pulA*^*+*^ strain RL09 cultivated on galactose; RL10 GLU, expression levels in *pulA*^*+*^ strain RL10 cultivated on glucose; RL10 GLY, expression levels in *pulA*^*+*^ strain RL10 cultivated on glycogen; ^2^LogR_1_, ^2^log ratio of expression levels of strains RL10 and RL09 both cultivated on galactose (fold change of expression calculated from LFQ-values from maxquant); ^2^logR_2_, ^2^log ratio of expression levels of *pulA*^*+*^ strain RL10 cultivated on glycogen and glucose.

| ACC NO | DESCRIPTION | PEPT | RL09 GAL | RL10 GAL | RL10 GLU | RL10 GLY | <sup>2</sup>logR<sub>1</sub> | <sup>2</sup>logR<sub>2</sub> |
| --- | --- | --- | --- | --- | --- | --- | --- | --- |
| A0A4Q0LMV5 | Alpha-dextrin endo-1,6-alpha-glucosidase | 64 | 2.82E+05 | 2.53E+07 | 7.49E+05 | 1.79E+07 | 6.49 | 4.58 |
| A0A135Z5X6 | Cell surface protein | 11 | 1.52E+06 | 3.17E+06 | 1.17E+06 | 3.18E+06 | 1.06 | 1.45 |
| D5GXS8 | Lysin(Glycoside Hydrolase Family 25) | 10 | 2.88E+05 | 3.65E+05 | 4.05E+05 | 9.74E+05 | 0.34 | 1.27 |
| D5H1Q3 | Aggregation promoting factor | 4 | ND | ND | 1.49E+05 | 2.22E+05 | ND | 0.57 |
| A0A135ZFI7 | N-acetylmuramidase | 17 | 1.30E+06 | 2.04E+06 | 3.99E+06 | 4.10E+06 | 0.65 | 0.04 |
| K1MET7 | Uncharacterized protein | 29 | 6.39E+06 | 1.02E+07 | 7.64E+06 | 7.85E+06 | 0.67 | 0.04 |
| A0A6M1GK10 | S-layer protein (Fragment) | 32 | 4.18E+05 | 6.86E+05 | 3.03E+05 | 2.99E+05 | 0.72 | -0.02 |
| C7XI68 | SlpA domain-containing protein | 14 | 3.20E+05 | 7.30E+05 | 6.40E+05 | 6.08E+05 | 1.19 | -0.07 |
| A0A6B8SPR2 | SlpA domain-containing protein | 15 | 3.08E+07 | 2.16E+07 | 1.29E+07 | 1.12E+07 | -0.51 | -0.20 |
| A0A135ZF64 | Amidase | 10 | 5.72E+05 | 1.70E+06 | 1.69E+06 | 1.43E+06 | 1.57 | -0.24 |
| A0A135ZH73 | Lysin | 11 | 4.52E+06 | 8.78E+06 | 6.58E+06 | 5.58E+06 | 0.96 | -0.24 |
| A0A7V8KTK5 | Putative surface layer protein | 48 | 1.57E+07 | 2.83E+07 | 2.68E+07 | 2.24E+07 | 0.85 | -0.26 |
| E3R4X6 | Uncharacterized protein | 9 | 2.58E+06 | 2.84E+06 | 2.73E+06 | 2.28E+06 | 0.14 | -0.26 |
| K1MJV3 | Uncharacterized protein | 22 | 1.64E+07 | 2.89E+07 | 2.03E+07 | 1.61E+07 | 0.82 | -0.33 |
| V5EI62 | Lysin (Glycoside Hydrolase Family 25) | 15 | 2.41E+06 | 4.60E+06 | 1.06E+07 | 7.62E+06 | 0.93 | -0.48 |
| V5EF53 | SlpA domain-containing protein | 12 | 2.09E+07 | 3.00E+07 | 6.12E+07 | 4.35E+07 | 0.52 | -0.49 |
| A0A4Q0LUE2 | Uncharacterized protein | 24 | 2.73E+07 | 1.81E+07 | 3.31E+07 | 2.32E+07 | -0.59 | -0.52 |
| A0A4R6CSS1 | Cell surface protein (SlpA) | 21 | 2.41E+06 | 5.85E+06 | 6.65E+06 | 4.56E+06 | 1.28 | -0.54 |
| D0DJ09 | Bacterial surface layer protein (Fragment) | 45 | 3.57E+08 | 4.85E+08 | 5.15E+08 | 3.51E+08 | 0.44 | -0.55 |
| K1M186 | Uncharacterized protein | 12 | 7.96E+06 | 9.68E+06 | 9.50E+06 | 6.14E+06 | 0.28 | -0.63 |
| A0A135YZ62 | Bacterial surface layer protein (SlpA) | 32 | 1.96E+07 | 3.35E+07 | 8.67E+07 | 5.52E+07 | 0.78 | -0.65 |
| D0DIV9 | Uncharacterized protein | 36 | 5.32E+07 | 7.57E+07 | 1.32E+08 | 7.87E+07 | 0.51 | -0.75 |
| V5GA31 | Lactocepin s-layer protein | 10 | 1.76E+06 | 2.75E+06 | 8.36E+06 | 3.22E+06 | 0.64 | -1.38 |
| A0A120DHV3 | Uncharacterized protein | 16 | 6.03E+06 | 1.07E+07 | 2.06E+07 | 7.70E+06 | 0.82 | -1.42 |
| K1N369 | Uncharacterized protein | 8 | 9.80E+04 | 3.70E+04 | 1.39E+05 | 3.01E+04 | -1.41 | -2.21 |
| A0A125P6L8 | S-layer protein (SlpA) | 13 | 1.09E+04 | 1.28E+05 | 8.69E+05 | 4.24E+03 | 3.55 | -7.68 |
| A0A7T1TNA6 | Bacterial surface layer protein | 30 | 4.71E+06 | 5.71E+06 | 6.07E+07 | 2.49E+05 | 0.28 | -7.93 |
| A0A1C2D5C0 | Cell surface protein | 5 | ND | 1.48E+06 | 8.12E+05 | ND | ND | ND |
| fraction of total detected proteins (iBAQ) |  |  | 0.47 | 0.46 | 0.22 | 0.40 |  |  |

### 2.4 Metagenome analysis

Our previous comparative genomics analysis of a collection of *Lactobacillus crispatus* strains isolated from the outpatient clinic in Amsterdam revealed variations in the N-terminal signal peptide region of the pullulanase gene that correlated with the ability of a strain to grow on glycogen [13]. In this strain collection, we found seven pullulanase-deficient strains sharing four different types of disruptions of the N-terminal sequence (Table 2). Although not all published *L. crispatus* genomes show the presence of a *pulA* gene [15], the N-terminal disruptions of the *pulA* sequence observed in our collection were not found in other *L. crispatus* genomes reported thus far.

**Table 2:** Overview of pullulanase variants in our strain collection of human vaginal *Lactobacillus crispatus* isolates [13].

| Strain | *pulA* variant |
| --- | --- |
| RL03, RL10, RL11, RL13, RL14, RL15, RL16, RL20, RL21, RL22, RL23, RL24, RL25, RL27, RL28, RL30, RL33 | Wild type |
| RL02 and RL09 | Two nucleotide deletion |
| RL06 and RL07 | N-terminal insertion of mobile element (variant 1) |
| RL19 and RL26 | N-terminal insertion of mobile element (variant 2) |
| RL05 | Large deletion |

We hypothesize that the isolation of these *pulA* variants is not a result of selective enrichment during the isolation procedure, but that *Lactobacillus crispatus* strains with defective pullulanase genes are common in the vaginal microbiota. To test this hypothesis we assessed the prevalence and frequency of disruptive *pulA* gene sequence variants such as those observed in our strain collection (Table 2). We analysed vaginal metagenome datasets from 1507 vaginal microbiota samples [28]. To our knowledge, this is the largest publicly available vaginal microbiome data collection which includes samples from several different studies of North American, African, and Chinese women. In total, we analysed 28.6 billion sequencing reads that made up 3.9 tera base pairs (Tbp) of metagenome sequencing data (on average, 19 million reads and 2.6 Gbp data per sample).

We first determined the prevalence of small but disruptive variations across the *pulA* gene sequence, similar to those observed in *pulA*^*-*^ strains RL02 and RL09 (Table 2). Out of 1507 samples we aligned to the *L. crispatus* RL10 *pulA* gene sequence, 270 resulted in alignments with a minimum of 10 times full length coverage. Variant calling on these 270 samples resulted in a total of 3202 unique variants (Appendix Figure A3). After low-frequency (allele frequency lower than 0.05) and low prevalence (observed in less than 3 samples) variants were filtered out, 54 variants remained, of which 19, 28 and 6 were low (e.g. synonymous mutations), moderate (*e.g*. mutations that cause an amino acid change), and high impact (stop gain or frameshift mutations), respectively (Figure 4).

**Fig. 4.**
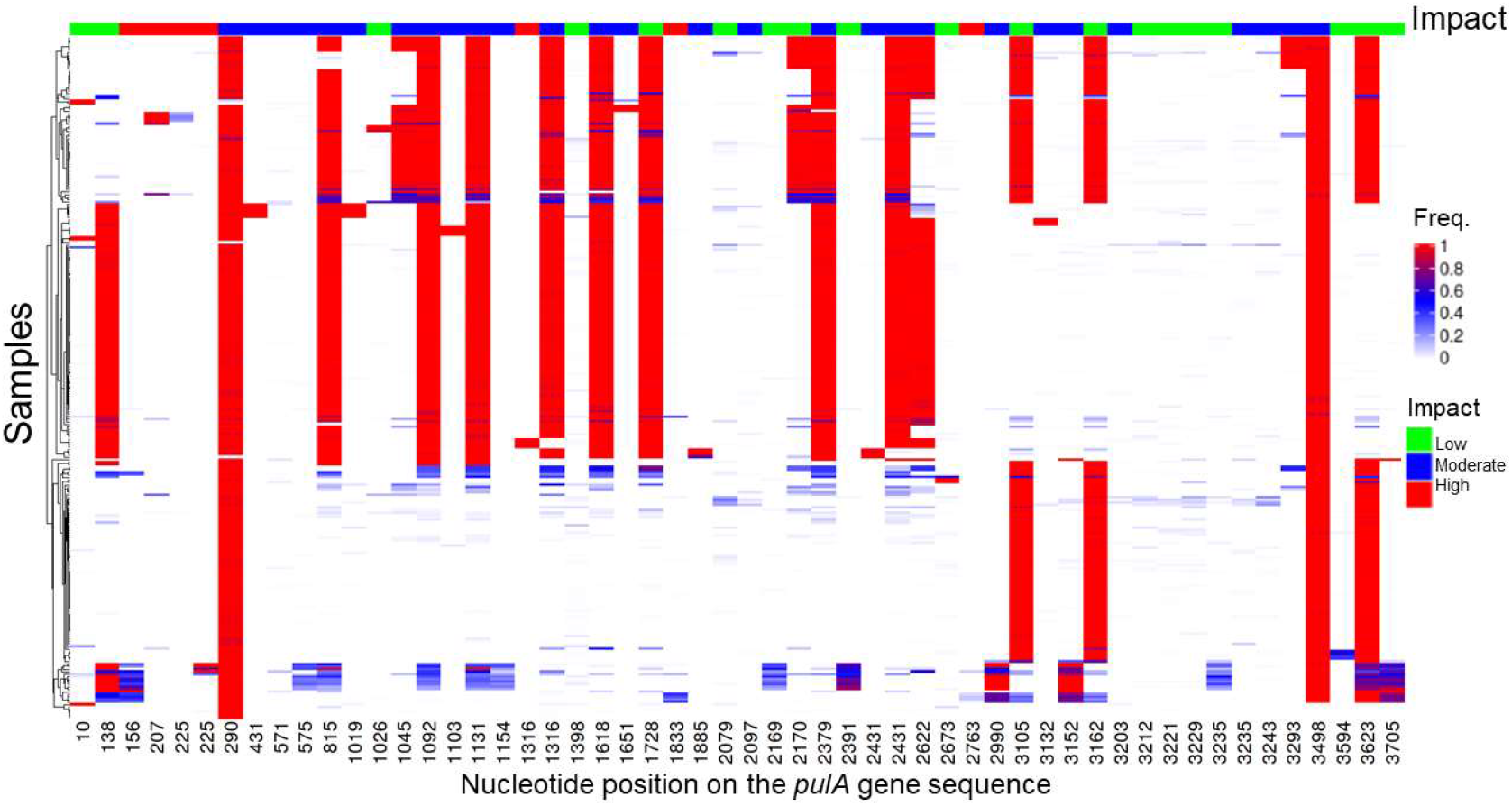
Small variants in the *Lactobacillus crispatus* pullulanase gene in the metagenome of vaginal microbial communities. The list and frequency of SNPs and short indels (small variants) identified in the 270 vaginal microbiome samples that showed at least ten times full length coverage for the *pulA* gene sequence. The variants were classified in terms of their impact on the protein sequence.

In 15 out of 270 samples (5.5%), at least one of the high impact mutations was observed at a high allele frequency (>0.7), where we expect a pullulanase deficient phenotype (Supplementary file pulA_small_variants.xlsx). Three of these high-impact mutations were near the N-terminus of the *pulA* sequence at positions 156, 207, and 225, impacting 11 samples in total.

Subsequently, we analysed the same set of 270 samples for larger structural variants compared to the pulA^+^ RL10 *pulA* gene sequence. In 56 out of 270 samples (20%), we identified large structural variants which had a high frequency (>0.7) (Supplemental File pulA_structural_variants.xlsx). The disruptive structural variation in 17 out of 56 of these samples was found be due to the insertion of transposase sequences into the pullulanase gene sequence, as in the case of the *pulA* locus from strains RL06, RL07, RL19, and RL26 in our own strain collection. Taken together, we identified high-frequency mutations in 62 of 270 *L. crispatus pulA* strains indicative of a dysfunctional pullulanase of the dominant strain and the inability of *L. crispatus* to contribute to the acidification of the vaginal environment with glycogen as a carbon and energy source.

## 3. Discussion

In this study we focus on genetic variation and disruption of the *Lactobacillus crispatus* pullulanase gene. Pullulanase is thought to play a central role in vaginal metabolomics. It catalyzes the first step of a metabolic pathway turning the most abundant vaginal carbohydrate (glycogen) into the most abundant metabolite (lactate) in one of the most abundant vaginal species *Lactobacillus crispatus*. In a previous study we published 33 genomes of vaginal *Lactobacillus crispatus* isolates that displayed remarkable variation in the gene locus. Strains with a disrupted pullulanase gene, through small indels or structural variants, were incapable of growth on glycogen.

The wealth of publically available metagenome datasets, repositories and gene catalogs allows for a more hypothesis driven research approach concentrating on specific genes that are associated with *in vivo* functionality. In the largest available metagenome database a substantial number of samples had similar genetic variation as were present in our previously reported whole genome sequences of *Lactobacillus crispatus* isolates. These variations consisted of both small deletions as well as relatively large structural variations disrupting the functionality of this gene.

Disruptions in the secretome of catalytically active enzymes are a well-known phenomenon in the microbial world. Bacteria may take advantage of cells in close proximity that cleave ‘public goods’ such as proteins or glycans into smaller peptides or maltodextrins. For instance, a protease-positive *Lactococcus lactis* population gradually lost protease activity in the entire population in a spatial-dependent manner. In this case the “cheating” proteolytic negative strains can utilize peptides supplied by the proteolytic positive strains without having the cost of protease production [29]. These disruptions may provide a fitness advantage to isolates, especially when, as we show in our mass spectrometric analysis, the enzyme forms a substantial fraction of the outer surface layer.

In the metagenome data of samples we did not find evidence of extensive variety of pullulanase sequences within a single vaginal microbiome: most samples only showed presence of one pullulanase variant, which we assume coincides to the dominant presence of one strain. We hypothesize that the metabolic advantage that a cheating strain in complex ecological samples may experience is not so much a matter of *intra*species competition, but rather due to *inter*species crossfeeding of metabolites. The presence of other glycogen degrading bacteria or human amylase may create an environment in which a *Lactobacillus crispatus* strain may obtain energetic advantages from shutting down the expression and secretion of S-layer associated enzymes, while benefiting from the breakdown products of extracellular glycogen hydrolysis by other species.

A recent study did not find such evolutionary pressure to mutate the pullulanase: even after cultivation of up to 1,000 generations by sequential sub-culturing in synthetic vaginal fluid, no genetic variations in the *pulA* locus of strains of *Lactobacillus crispatus* were detected. [30]. We hypothesize that this may be due to the available carbohydrates in this simulated vaginal fluid, which contains both glucose and glycogen and is further supplemented with amylase which degrades the glycogen into smaller maltodextrins. Our study implies that *L. crispatus* grown in such an environment will first grow on glucose, maltose and maltotriose and will not express pullulanase until these carbohydrates are depleted. This will diminish the advantage of cheating or any other evolutionary pressure on the production of alpha-glucosidase.

Genetic disruption affected a minority of vaginal samples in the metagenomic database or isolates in our strain collection. In most environments *L. crispatus* maintains an intact pullulanase gene with no sign of a genetic alteration disrupting its activity. However, our study shows that carbon catabolite repression may further reduce activity of *Lactobacillus crispatus* pullulanase activity *in vivo*. Bacteria generally regulate metabolic pathways optimizing their growth strategy by prioritizing carbon sources based on nutritional quality. The presence of glucose, maltose and maltotriose in the vagina has been well-documented, especially in *Lactobacillus*-dominated environments [31,32].

Given the prevalence of these carbon sources in metabolomics studies, as well as human amylase we expect that *Lactobacillus crispatus* in the vaginal environment preferentially relies on glycogen cleavage by other bacteria or host amylase and only expresses pullulanase when other carbon sources are depleted.

## 4. Materials and Methods

### 4.1 Standard cultivation conditions

*Lactobacillus crispatus* pullulanase deficient strain (*L. crispatus* RL09) and a pullulanase sufficient strain (*L. crispatus* RL10) were routinely cultivated on NYCIII medium with HEPES (2.4 g/L), proteose peptone #3 (Becton Dickinson, 211693) (15 g/L), yeast extract (Oxoid, LP0021) (3.8 g/L), NaCl (5 g/L), glucose monohydrate (Santa Cruz Biotechnology, 14431-43-7) (5.5 g/L g), heat inactivated horse serum (10%) and for agar plates 1,5% (w/v) agarose. Plates, precultures and subcultures were grown under anaerobic conditions (gas mixture of 10% CO_2_ and 90% N_2_ at 37°C.

### 4.2 PCR and sequencing of signal peptide region L. crispatus pullulanase

Previously, Illumina-whole genome sequencing showed a two nucleotide deletion in the N-terminal *pulA* signal peptide sequence of *Lactobacillus crispatus* RL09. In order to confirm this frame shift mutation we performed genetic analysis of the area with the deletion by PCR amplification and Sanger sequencing. Genomic DNA was isolated from a 24 hour liquid culture using the GenElute™ Bacterial Genomic DNA Kit (Merck). A 300 basepair sequence at the start site of the pullulanase was amplified with forward primer GCAAATGAAAGCGCATACGTTT (annealing 42 basepairs upstream the start site) and reverse primer TGTTGACGCTGCTTTGCTT (annealing 257 basepairs downstream the start site) using a High Fidelity Phusion® DNA polymerase (ThermoFisher). The size of the PCR product was verified with ethidium bromide on gel to be the predicted size of 299 bp. The PCR product was purified using the GeneJET PCR Purification Kit (Thermo Fisher) and sequenced by Sanger Dideoxy sequencing (Eurofins).

### 4.3 Serial propagation on various carbohydrates

Growth on various carbohydrate sources was tested with NYCIII without glucose supplemented with D-lactose monohydrate (Sigma-Aldrich), D-raffinose pentahydrate (Alfa Aesar), D-cellobiose (Sigma-Aldrich), D-melibiose (Sigma-Aldrich), D-maltose monohydrate (Fluka), D-trehalose dihydrate (Sigma-Aldrich), D-galactose (Sigma-Aldrich), lactulose (Sigma-Aldrich), glucose monohydrate (Merck), bovine glycogen (Sigma-Aldrich) and oyster glycogen (Alfa Aesar). Growth of *L. crispatus* RL09 and *L. crispatus* RL10 on these carbohydrate sources was examined by serial propagation on glucose-free NYCIII medium supplemented with 5% (w/v) of the different carbohydrates. The serial propagation was executed using two passages of a twentyfold sub-culturing in a 96-wells plate. Between each passage, the cells were anaerobically grown for 24h in jars (Oxoid). To monitor growth, optical density was measured at 600 nm in a sterile 96-well flat bottom plates in a spectrophotometer (Spectramax Plus 384; Molecular devices) after ten times dilution in sterile phosphate-buffered saline solution (PBS). Acidification of the medium by *L. crispatus* was assessed by pH measurements (HI 2210 pH meter, Hanna instrument) after every passage after 24h of anaerobic growth. Statistical testing of optical densities and pH values were done with t-tests: two samples assuming unequal variations (α = 0.05, two-sided).

### 4.4 The starch degradation assay

Glycogen is a polymer of α-1,4-linked and α-1,6-linked glucose residues. Starch degradation, measured in a basic starch-iodine assay, can be used as a proxy since the amylopectin moiety of starch has a similar branched structure with α-1,4-linked and α-1,6-linked glucose residues. To test α-glucosidase activity strains were grown for 24 hours. Cells were spun down and resuspended in amylase buffer (100 mM Na-Acetate with 5 mM CaCl_2_, pH 5,3). 50 uL supernatant or resuspended pellet was added to 150 uL of 10 g/L starch solution (Sigma, S9765) in amylase buffer. Chloramphenicol (Sigma, C0378-25G) from a stock solution in ethanol was added in a final concentration of 10 ng/mL to prevent growth. 50 μL of resuspended pellet of culture spent medium was mixed with 150 μL of this starch working solution and incubated at 37°C for 24 hours. To visualize residual starch, 10 uL was added to 290 uL of an iodine stock solution consisting of 0,2 g I2 and 2 g Kl in 100 mL 50 mM HCl. Absorption was measured at 600 nm in a 96 well plate reader. Experiments were carried out with a minimum of (two) biological replicates for RL09 on maltose and maltotriose; all other conditions had between 3 and 9 replicates carried out on separate days by a total of 4 independent experiments

### 4.5 Enrichment for surface layer (associated) proteins

The enrichment for SLP’s and SLAP’s was modified from a standard LiCl S-layer extraction protocol for *L. acidophilus* (Goh et al., 2009; Lortal et al., 1992). Bacterial cells of *L. crispatus* were grown to stationary phase (24 h), centrifuged at 2236 g for 10 min (4°C), and washed twice with 25 ml cold PBS (Gibco), pH 7.4. Cells were agitated for 15 min at 4°C following the addition of 5 M LiCl (Fisher Scientific). Supernatants, containing SLP’s and SLAP’s, were harvested via centrifugation at 8994 g for 10 min (4°C) and transferred to a 6000–8000 kDa Spectra/Por molecular porous membrane (Spectrum Laboratories) and dialysed against cold distilled water for 24 h, changing the water every 2 h for the first 8 h. The dialysed precipitate was harvested via centrifugation at 20,000 g for 30 min (4°C).

### 4.6 Sample Preparation for Proteomics Analysis

Protein pellets were resuspended in 200 μL of digestion buffer (5 mM DTT, 50 mM NH_4_HCO_3_, pH 8.0) and incubated at 55°C for one hour to reduce disulfide bridges. Then samples were treated with 15 mM iodoacetamide (IAA) to alkylate free cysteines in the dark for 45 min. Subsequently samples were digested by trypsin (1:100 w/w protease:protein ratio) at 37°C for 18 hours. The mixture of peptides was desalted using C18 solid phase extraction columns (Omix, Agilent) following the manufacturer’s protocol. The final peptide mixture was dried in a vacuum centrifuge and suspended in 0.1% formic acid in water (ULC-MS grade, Biosolve) for MS analysis.

### 4.7 LC-MS/MS Analysis

Mass spectrometric analysis of 200 ng peptides was carried out on a timsTOF Pro (Bruker, Bremen, Germany) equipped with an Ultimate 3000 nanoRSLC UHPLC system (Thermo Scientific, Germeringen, Germany). Samples were injected onto a C18 column (75 μm, 250 mm, 1.6 μm particle size, Aurora, Ionopticks, Fitzroy, Australia) kept at 50°C. Peptides were loaded at 400 nl/min for 1 minute in 3% solvent B and separated by a multi-step gradient: 5% solvent B at 2 min 17%, solvent B at 24 min, 25% solvent B at 29 min, 34% solvent B at 32 min, 99% solvent B at 33 min held untill 40 minute, returning to 3% solvent B at 40.1 min and held untill 58 min to recondition the column (Solvent A: 0.1% formic acid in water, Solvent B: 0.1% formic acid in acetonitrile). MS analysis of eluting peptides was performed by a time-of-flight mass spectrometer. The precursor scan ranged from 100 to 1700 m/z and a tims range of 0.6–1.6 V.s/cm2 in PASEF mode. A total of 10 PASEF MS/MS scans following collision induced dissociation (collision energy from 20–59 eV) were collected with a total cycle time of 1.16 s.

Raw MS/MS data were processed using Maxquant software (version 1.6.14.0) [33] earching a pan proteome database of *L. crispatus* (Uniprot, downloaded 1-11-2021). To estimate false spectrum assignment rate a reverse version of the same database was also searched. The settings were as follows: Enzyme Trypsin/P allowing for a maximum of two missed cleavages, variable modifications: Oxidation (M), fixed modifications: Carbamidomethyl (C). Settings were default for timsDDA, match between runs was enabled with a matching time window of 0.2 minutes and a matching ion mobility window of 0.05 indices. For label free quantification, both iBAQ and LFQ were enabled [33].

### 4.8 Analysis of the pulA gene region

The contig of *Lactobacillus crispatus* RL10 was analyzed with SnapGene version 5.3.2. (GSL Biotech LLC). The promoter sequences (−35 and -10 elements) were predicted by use of BPROM (Prediction of Bacterial Promoters) and FindTerm (Finding Terminators in bacterial genomes) [34]. The cabolite responsive element (*cre* site) has been identified by matching the palindromic nucleotide motif of *Lactobacillus plantarum* (TGWAANCGNTNWCA) [35]

### 4.9 Analysis of publicly available vaginal metagenomes

The raw shotgun metagenome sequencing data of 1507 publicly available vaginal microbiome samples were downloaded from the VIRGO database [28]. In addition to the 211 samples which were sequenced for the construction of VIRGO, this large data collection contains samples from other studies of North American [36], African [5], and Chinese women [37]. The Illumina reads were processed using FastQC to determine data statistics such as data yield per sample. The samples were taxonomically characterised using Kraken 2 [38] and Bracken [39]. The human read counts in the species-level results from Bracken were removed and the data was normalised to obtain relative abundances of non-human taxa such as bacteria, archaea, viruses, and fungi. Taxa that had less than 0.5% relative abundance and which were observed in less than 5% of the samples were filtered out. The downstream taxonomic analyses and visualisation were performed using R v3.6.3 (R Core Team, 2020) with R packages vegan, phyloseq [40], ComplexHeatmap [41], and ggplot2 v3.3.3.

### 4.10 Pullulanase sequence variant detection

The raw reads were aligned to the RL10 *pulA* gene sequence using bowtie2 [42] with default parameters except for “--very-sensitive-local --no-unal”. Alignment statistics for each sample such as coverage were calculated using samtools v1.13 [43]. Single-nucleotide polymorphisms (SNPs) and short indels in samples with a minimum 10X coverage were determined using FreeBayes v1.2 [44] in a multi-sample variant calling approach using default parameters except for “–min-alternate-count 2 –min-coverage 10 –min-alternate-fraction 0.01 –use-duplicate-reads –min-mapping-quality 0 –ploidy 1 –pooled-continuous”.

The variants were annotated in terms of their impact on the RL10 *pulA* protein sequence using SnpEff [45]: high (*e.g*. mutations that result in frameshifts or stop gain), moderate (e.g. mutations that result in an amino acid change), and low (synonymous mutations). The variant allele frequency heatmap for variants which had at least 0.05 alternate allele frequency (i.e., supported by at least 5% of the reads at that position) and were observed in at least 3 samples was plotted using R packages VariantAnnotation [46], and ggplot2. SNPs and short indels with a minimum frequency of 0.7 were classified as high-frequency short variants.

The same sample set used in small variant detection which had a minimum coverage of 10 times along the RL10 *pulA* sequence was used for detecting larger and structural variations such as insertions, deletions, inversions, translocations, and duplications in read mappings. To this end, the indels and Structural Variants tool in CLC Genomics v.12.0.3 was used with default parameters (P-value threshold=0.0001) except for “minimum number of reads=10”. Structural variants with a minimum frequency of 0.7 were classified as high-frequency large variants.

### 4.11 Identification of pulA genes with transposase insertions

To create a database of *L. crispatus* transposase gene sequences, we first downloaded 147 annotated *L. crispatus* genomes from NCBI (assembly level: scaffold, chromosome, and complete). We then created a FASTA file of 19,551 redundant transposase gene sequences by combining the coding sequences which had a ‘transposase’ annotation in the 147 *L. crispatus* genomes. This database was clustered at 95% sequence identity using CD-HIT using default parameters [47] to obtain a non-redundant transposase database of 508 gene sequences.

To identify *pulA*-rich samples that had transposase insertions, we mapped the reads which were mapped to the *L. crispatus* RL10 *pulA* sequence to the above-mentioned transposase database using bowite2 as described above. The list of samples where more than 20 transposase reads were found was compared to the list of samples with high-frequency structural variants to determine the impact of transposase insertions in causing disruptive structural variations.

## Supplementary Materials

The following are available online at www.mdpi.com/xxx/s1,

Supplementary file data Figure 1

Supplementary file data Figure 2 and Figure A2

Supplementary file pulA_small_variants.xlsx

Supplementary file pulA_structural_variants.xlsx

## Author Contributions

Conceptualization, RYH, RK; Methodology, RYH, GK, RK; Software, AM; Experiments: RYH, GK, IvV, DM, RK; data curation: AM, RYH, Writing—original draft preparation, RYH, AM, GK, RK; Writing – review and editing RYH, RK; Supervision, RYH, DM, RK; All authors have read and agreed to the published version of the manuscript.”

## Institutional Review Board Statement

Not applicable

## Informed Consent Statement

Not applicable

## Data Availability Statement

All data presented here are included in the manuscript, appendix or supplemental file. The mass-spectrometry data have been deposited in Proteome Exchange.

## Acknowledgments

We would like to thank the MSc-students Omar Beganovic, Deborah Jekel, Yumiko van Diest, Leon Steenbergen, and Ritesh Panchoe for their contributions to the research presented in this publication.

## Appendix A

The appendix is an optional section that can contain details and data supplemental to the main text—for example, explanations of experimental details that would disrupt the flow of the main text but nonetheless remain crucial to understanding and reproducing the research shown; figures of replicates for experiments of which representative data is shown in the main text can be added here if brief, or as Supplementary data. Mathematical proofs of results not central to the paper can be added as an appendix.

## Appendix A

**Figure A1.**
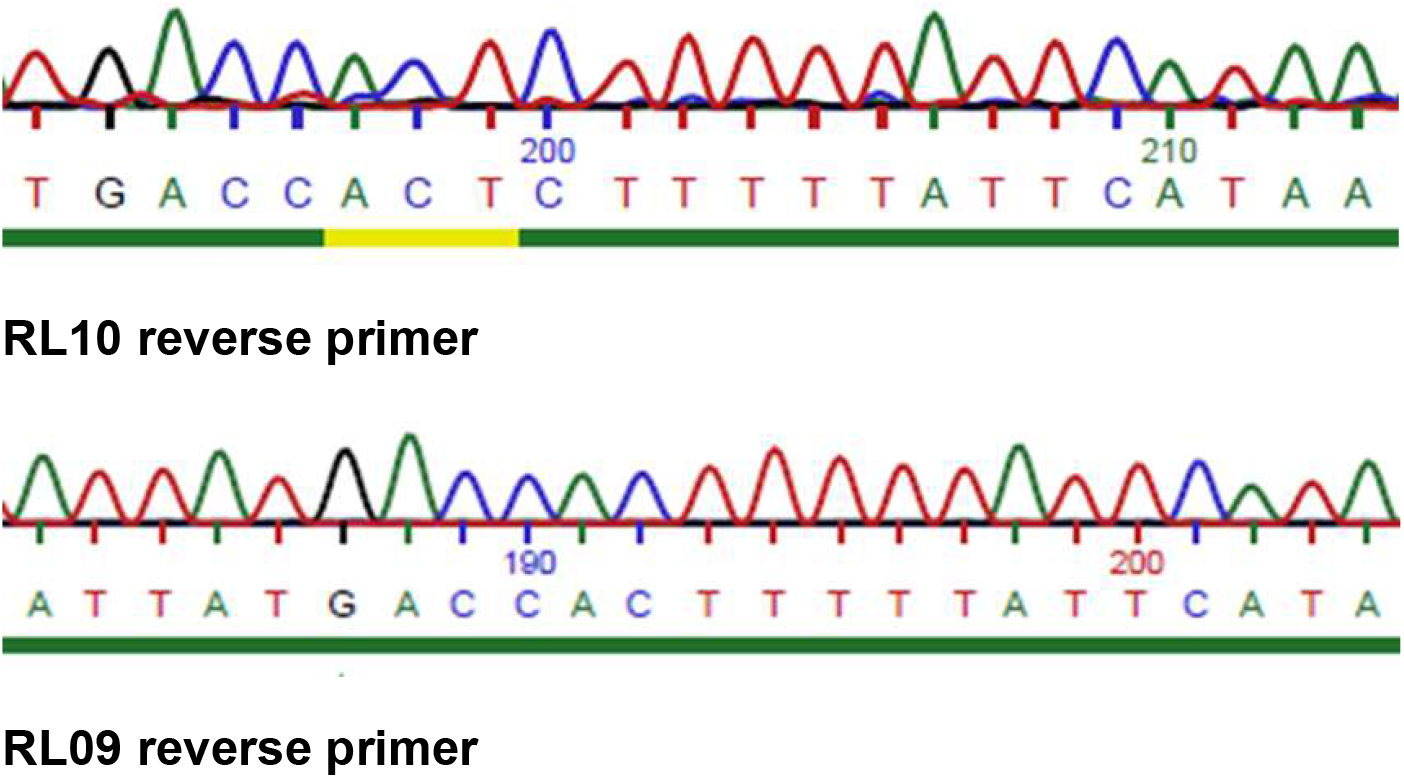
Resequencing the *Lactobacillus crispatus pulA* signal peptide region of a *pulA*^*+*^ strains RL10 and RL09.

**Figure A2.**
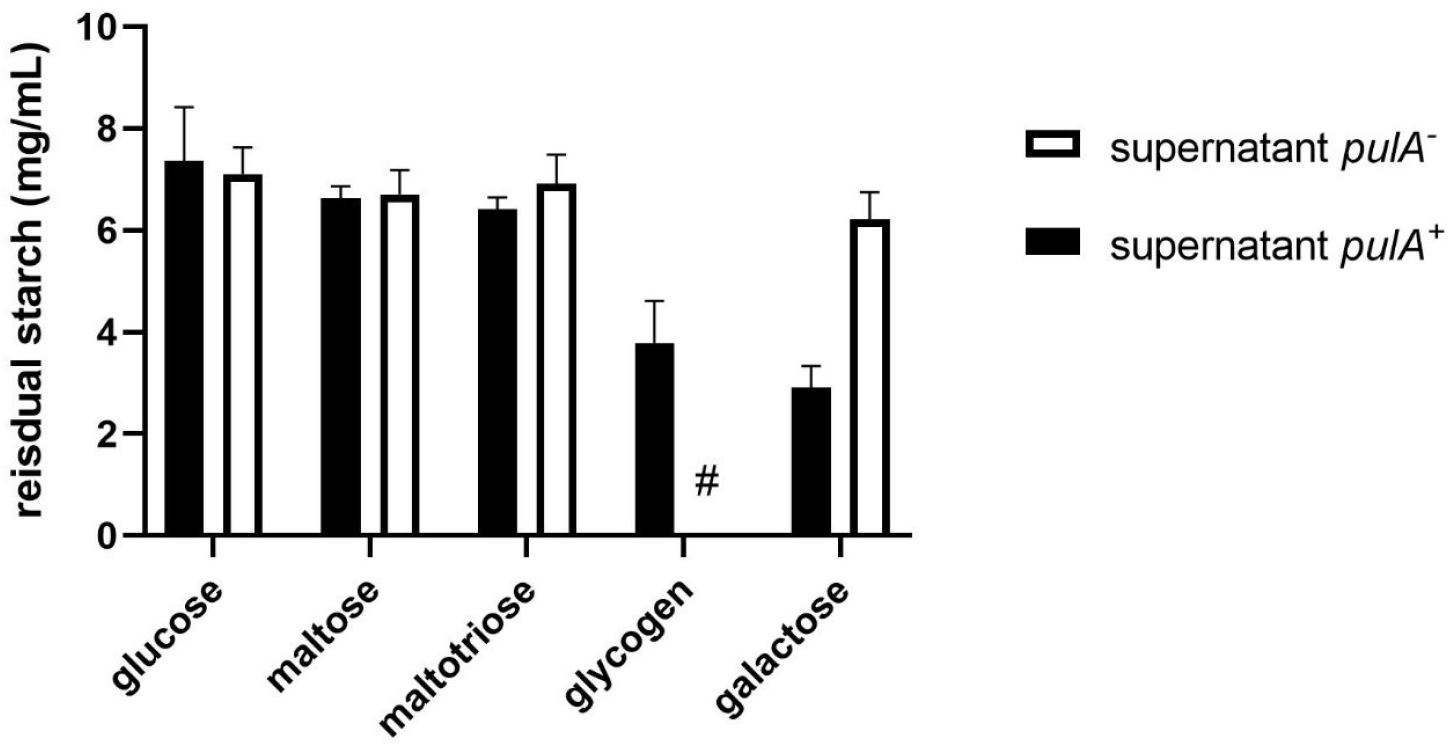
Starch degrading activity in spent medium of *Lactobacillus* crispatus pullulanase sufficient and deficient strains grown on NYC-, glucose, maltose, maltotriose, glycogen (only *pulA*^*+*^) and galactose. Cells were cultured in NYCIII medium supplemented with these carbohydrates and after centrifugation and resuspension of the pellet incubated in a starch solution (7.5 g/L). Experiments were carried out with a minimum of (two) biological replicates; averages and standard deviations are shown. Asterisks indicate *p*-values calculated by Mann Whitney of * p<0,05, ** p<0,005, *** p<0,0005.

**Figure A3.**
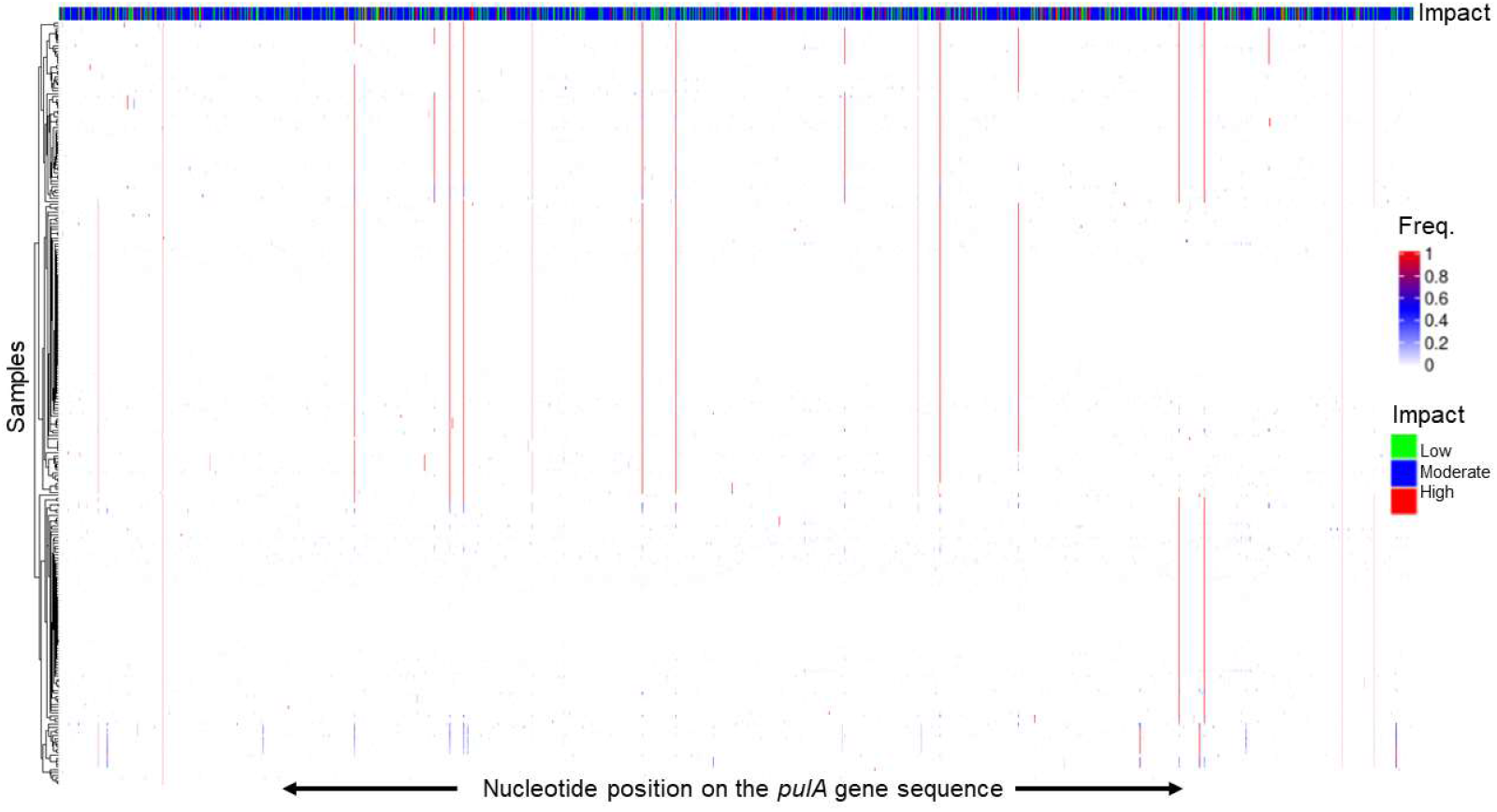
Variant calling on the *pulA* gene positive 270 samples in the vaginal metagenome data set.

## References

1. O’Hanlon, D.E.; Moench, T.R.; Cone, R.A. Vaginal PH and Microbicidal Lactic Acid When Lactobacilli Dominate the Microbiota. PloS One 2013, 8, e80074, doi:10.1371/journal.pone.0080074.

2. Mirmonsef, P.; Hotton, A.L.; Gilbert, D.; Burgad, D.; Landay, A.; Weber, K.M.; Cohen, M.; Ravel, J.; Spear, G.T. Free Glycogen in Vaginal Fluids Is Associated with Lactobacillus Colonization and Low Vaginal PH. PloS One 2014, 9, e102467, doi:10.1371/journal.pone.0102467.

3. Ravel, J.; Gajer, P.; Abdo, Z.; Schneider, G.M.; Koenig, S.S.; McCulle, S.L.; Karlebach, S.; Gorle, R.; Russell, J.; Tacket, C.O.; et al. Vaginal Microbiome of Reproductive-Age Women. Proc. Natl. Acad. Sci. U. S. A. 2011, 108 Suppl 1, 4680–4687, doi:10.1073/pnas.1002611107;

4. Dols, J.A.M.; Molenaar, D.; van der Helm, J.J.; Caspers, M.P.M.; de Kat Angelino-Bart, A.; Schuren, F.H.J.; Speksnijder, A.G.C.L.; Westerhoff, H.V.; Richardus, J.H.; Boon, M.E.; et al. Molecular Assessment of Bacterial Vaginosis by Lactobacillus Abundance and Species Diversity. BMC Infect. Dis. 2016, 16, doi:10.1186/s12879-016-1513-3.

5. Gosmann, C.; Anahtar, M.N.; Handley, S.A.; Farcasanu, M.; Abu-Ali, G.; Bowman, B.A.; Padavattan, N.; Desai, C.; Droit, L.; Moodley, A.; et al. Lactobacillus-Deficient Cervicovaginal Bacterial Communities Are Associated with Increased HIV Acquisition in Young South African Women. Immunity 2017, 46, 29–37, doi:10.1016/j.immuni.2016.12.013.

6. De Seta, F.; Lonnee-Hoffmann, R.; Campisciano, G.; Comar, M.; Verstraelen, H.; Vieira-Baptista, P.; Ventolini, G.; Lev-Sagie, A. The Vaginal Microbiome: III. The Vaginal Microbiome in Various Urogenital Disorders. J. Low. Genit. Tract Dis. 2022, 26, 85–92, doi:10.1097/LGT.0000000000000645.

7. Ceccarani, C.; Foschi, C.; Parolin, C.; D’Antuono, A.; Gaspari, V.; Consolandi, C.; Laghi, L.; Camboni, T.; Vitali, B.; Severgnini, M.; et al. Diversity of Vaginal Microbiome and Metabolome during Genital Infections. Sci. Rep. 2019, 9, 14095, doi:10.1038/s41598-019-50410-x.

8. Koedooder, R.; Singer, M.; Schoenmakers, S.; Savelkoul, P.H.M.; Morré, S.A.; de Jonge, J.D.; Poort, L.; Cuypers, W.; Beckers, N.G.M.; Broekmans, F.J.M.; et al. The Vaginal Microbiome as a Predictor for Outcome of in Vitro Fertilization with or without Intracytoplasmic Sperm Injection: A Prospective Study. Hum Reprod 2019, 34, 1042–1054, doi:10.1093/humrep/dez065.

9. Vergaro, P.; Tiscornia, G.; Barragán, M.; García, D.; Rodriguez, A.; Santaló, J.; Vassena, R. Vaginal Microbiota Profile at the Time of Embryo Transfer Does Not Affect Live Birth Rate in IVF Cycles with Donated Oocytes. Reprod. Biomed. Online 2019, 38, 883–891, doi:10.1016/j.rbmo.2018.12.019.

10. Fettweis, J.M.; Serrano, M.G.; Brooks, J.P.; Edwards, D.J.; Girerd, P.H.; Parikh, H.I.; Huang, B.; Arodz, T.J.; Edupuganti, L.; Glascock, A.L.; et al. The Vaginal Microbiome and Preterm Birth. Nat. Med. 2019, 25, 1012–1021, doi:10.1038/s41591-019-0450-2.

11. Usyk, M.; Schlecht, N.F.; Pickering, S.; Williams, L.; Sollecito, C.C.; Gradissimo, A.; Porras, C.; Safaeian, M.; Pinto, L.; Herrero, R.; et al. MolBV Reveals Immune Landscape of Bacterial Vaginosis and Predicts Human Papillomavirus Infection Natural History. Nat. Commun. 2022, 13, 233, doi:10.1038/s41467-021-27628-3.

12. Dols, J.A.M.; Reid, G.; Kort, R.; Schuren, F.H.J.; Tempelman, H.; Bontekoe, T.R.; Korporaal, H.; Van der Veer, E.M.; Smit, P.W.; Boon, M.E. PCR-Based Identification of Eight Lactobacillus Species and 18 Hr-HPV Genotypes in Fixed Cervical Samples of South African Women at Risk of HIV and BV. Diagn. Cytopathol. 2012, 40, 472–477, doi:10.1002/dc.21786.

13. van der Veer, C.; Hertzberger, R.Y.; Bruisten, S.M.; Tytgat, H.L.P.; Swanenburg, J.; de Kat Angelino-Bart, A.; Schuren, F.; Molenaar, D.; Reid, G.; de Vries, H.; et al. Comparative Genomics of Human Lactobacillus Crispatus Isolates Reveals Genes for Glycosylation and Glycogen Degradation: Implications for in Vivo Dominance of the Vaginal Microbiota. Microbiome 2019, 7, 49, doi:10.1186/s40168-019-0667-9.

14. Woolston, B.M.; Jenkins, D.J.; Hood-Pishchany, M.I.; Nahoum, S.R.; Balskus, E.P. Characterization of Vaginal Microbial Enzymes Identifies Amylopullulanases That Support Growth of Lactobacillus Crispatus on Glycogen; 2021; p. 2021.07.19.452977;

15. Pan, M.; Hidalgo-Cantabrana, C.; Barrangou, R. Host and Body Site-Specific Adaptation of Lactobacillus Crispatus Genomes. NAR Genomics Bioinforma. 2020, 2, lqaa001, doi:10.1093/nargab/lqaa001.

16. Nasioudis, D.; Beghini, J.; Bongiovanni, A.M.; Giraldo, P.C.; Linhares, I.M.; Witkin, S.S. Alpha-Amylase in Vaginal Fluid: Association With Conditions Favorable to Dominance of Lactobacillus. Reprod. Sci. Thousand Oaks Calif 2015, doi:1933719115581000.

17. Spear, G.T.; French, A.L.; Gilbert, D.; Zariffard, M.R.; Mirmonsef, P.; Sullivan, T.H.; Spear, W.W.; Landay, A.; Micci, S.; Lee, B.H.; et al. Human Alpha-Amylase Present in Lower-Genital-Tract Mucosal Fluid Processes Glycogen to Support Vaginal Colonization by Lactobacillus. J. Infect. Dis. 2014, 210, 1019–1028, doi:10.1093/infdis/jiu231.

18. Nunn Kenetta L.; Clair Geremy C.; Adkins Joshua N.; Engbrecht Kristin; Fillmore Thomas; Forney Larry J.; Young Vincent B. Amylases in the Human Vagina. mSphere 2020, 5, e00943–20, doi:10.1128/mSphere.00943-20.

19. Spear, G.T.; McKenna, M.; Landay, A.L.; Makinde, H.; Hamaker, B.; French, A.L.; Lee, B.H. Effect of PH on Cleavage of Glycogen by Vaginal Enzymes. PloS One 2015, 10, e0132646, doi:10.1371/journal.pone.0132646.

20. Collins, S.L.; McMillan, A.; Seney, S.; van der Veer, C.; Kort, R.; Sumarah, M.W.; Reid, G. Promising Prebiotic Candidate Established by Evaluation of Lactitol, Lactulose, Raffinose, and Oligofructose for Maintenance of a Lactobacillus-Dominated Vaginal Microbiota. Appl. Environ. Microbiol. 2018, 84, e02200–17, doi:10.1128/AEM.02200-17.

21. Møller, M.S.; Goh, Y.J.; Rasmussen, K.B.; Cypryk, W.; Celebioglu, H.U.; Klaenhammer, T.R.; Svensson, B.; Abou Hachem, M. An Extracellular Cell-Attached Pullulanase Confers Branched α-Glucan Utilization in Human Gut Lactobacillus Acidophilus. Appl. Environ. Microbiol. 2017, 83, e00402–17, doi:10.1128/AEM.00402-17.

22. Muscariello, L.; Marasco, R.; De Felice, M.; Sacco, M. The Functional CcpA Gene Is Required for Carbon Catabolite Repression in Lactobacillus Plantarum. Appl. Environ. Microbiol. 2001, 67, 2903–2907, doi:10.1128/AEM.67.7.2903-2907.2001.

23. Mahr, K.; Hillen, W.; Titgemeyer, F. Carbon Catabolite Repression in Lactobacillus Pentosus: Analysis of the CcpA Region. Appl. Environ. Microbiol. 2000, 66, 277–283.

24. Barrangou, R.; Azcarate-Peril, M.A.; Duong, T.; Conners, S.B.; Kelly, R.M.; Klaenhammer, T.R. Global Analysis of Carbohydrate Utilization by Lactobacillus Acidophilus Using CDNA Microarrays. Proc. Natl. Acad. Sci. U. S. A. 2006, 103, 3816–3821, doi:10.1073/pnas.0511287103.

25. Thompson, J.; Turner, K.W.; Thomas, T.D. Catabolite Inhibition and Sequential Metabolism of Sugars by Streptococcus Lactis. J. Bacteriol. 1978, 133, 1163–1174.

26. Qian, N.; Stanley, G.A.; Bunte, A.; Rdstrm, P. Product Formation and Phosphoglucomutase Activities in Lactococcus Lactis: Cloning and Characterization of a Novel Phosphoglucomutase Gene. Microbiol. Read. Engl. 1997, 143 (Pt 3), 855–865, doi:10.1099/00221287-143-3-855.

27. Åvall-Jääskeläinen, S.; Palva, A. Lactobacillus Surface Layers and Their Applications. FEMS Microbiol. Rev. 2005, 29, 511–529, doi:10.1016/j.fmrre.2005.04.003.

28. Ma, B.; France, M.T.; Crabtree, J.; Holm, J.B.; Humphrys, M.S.; Brotman, R.M.; Ravel, J. A Comprehensive Non-Redundant Gene Catalog Reveals Extensive within-Community Intraspecies Diversity in the Human Vagina. Nat. Commun. 2020, 11, 1–13, doi:10.1038/s41467-020-14677-3.

29. Bachmann, H.; Molenaar, D.; Kleerebezem, M.; van Hylckama Vlieg, J.E. High Local Substrate Availability Stabilizes a Cooperative Trait. ISME J. 2011, 5, 929–932, doi:10.1038/ismej.2010.179.

30. Brandt, K.; Barrangou, R. Adaptive Response to Iterative Passages of Five Lactobacillus Species in Simulated Vaginal Fluid. BMC Microbiol. 2020, 20, 339, doi:10.1186/s12866-020-02027-8.

31. Srinivasan, S.; Morgan, M.T.; Fiedler, T.L.; Djukovic, D.; Hoffman, N.G.; Raftery, D.; Marrazzo, J.M.; Fredricks, D.N. Metabolic Signatures of Bacterial Vaginosis. mBio 2015, 6, e00204–15, doi:10.1128/mBio.00204-15.

32. Gregoire, A.T. Carbohydrates of Human Vaginal Tissue. Nature 1963, 198, 996, doi:10.1038/198996a0.

33. MaxQuant Enables High Peptide Identification Rates, Individualized p.p.b.-Range Mass Accuracies and Proteome-Wide Protein Quantification | Nature Biotechnology Available online: https://www.nature.com/articles/nbt.1511 (accessed on 15 April 2022).

34. Solovyev, V.; Salamov, A. Automatic Annotation of Microbial Genomes and Metagenomic Sequences 3 MATERIAL AND METHODS Learning Parameters and Prediction of Protein-Coding Genes Available online: https://www.semanticscholar.org/paper/Automatic-Annotation-of-Microbial-Genomes-and-3-AND-Solovyev-Salamov/88cd5fdbfb2dba16a6b23987d30ad9437ff0c805 (accessed on 15 April 2022).

35. Lu, Y.; Song, S.; Tian, H.; Yu, H.; Zhao, J.; Chen, C. Functional Analysis of the Role of CcpA in Lactobacillus Plantarum Grown on Fructooligosaccharides or Glucose: A Transcriptomic Perspective. Microb. Cell Factories 2018, 17, 201, doi:10.1186/s12934-018-1050-4.

36. Human Microbiome Project Consortium A Framework for Human Microbiome Research. Nature 2012, 486, 215–221, doi:10.1038/nature11209.

37. Li, F.; Chen, C.; Wei, W.; Wang, Z.; Dai, J.; Hao, L.; Song, L.; Zhang, X.; Zeng, L.; Du, H.; et al. The Metagenome of the Female Upper Reproductive Tract. GigaScience 2018, 7, doi:10.1093/gigascience/giy107.

38. Wood, D.E.; Lu, J.; Langmead, B. Improved Metagenomic Analysis with Kraken 2. Genome Biol. 2019, 20, 257, doi:10.1186/s13059-019-1891-0.

39. Lu, J.; Breitwieser, F.P.; Thielen, P.; Salzberg, S.L. Bracken: Estimating Species Abundance in Metagenomics Data. PeerJ Comput. Sci. 2017, 3, e104, doi:10.7717/peerj-cs.104.

40. McMurdie, P.J.; Holmes, S. Phyloseq: A Bioconductor Package for Handling and Analysis of High-Throughput Phylogenetic Sequence Data. Pac. Symp. Biocomput. Pac. Symp. Biocomput. 2012, 235–246.

41. Gu, Z.; Eils, R.; Schlesner, M. Complex Heatmaps Reveal Patterns and Correlations in Multidimensional Genomic Data. Bioinforma. Oxf. Engl. 2016, 32, 2847–2849, doi:10.1093/bioinformatics/btw313.

42. Langmead, B.; Salzberg, S.L. Fast Gapped-Read Alignment with Bowtie 2. Nat. Methods 2012, 9, 357–359, doi:10.1038/nmeth.1923.

43. Li, H.; Handsaker, B.; Wysoker, A.; Fennell, T.; Ruan, J.; Homer, N.; Marth, G.; Abecasis, G.; Durbin, R.; 1000 Genome Project Data Processing Subgroup The Sequence Alignment/Map Format and SAMtools. Bioinforma. Oxf. Engl. 2009, 25, 2078–2079, doi:10.1093/bioinformatics/btp352.

44. Garrison, E.; Marth, G. Haplotype-Based Variant Detection from Short-Read Sequencing. 12073907 Q-Bio 2012.

45. Cingolani, P.; Platts, A.; Wang, L.L.; Coon, M.; Nguyen, T.; Wang, L.; Land, S.J.; Lu, X.; Ruden, D.M. A Program for Annotating and Predicting the Effects of Single Nucleotide Polymorphisms, SnpEff: SNPs in the Genome of Drosophila Melanogaster Strain W1118; Iso-2; Iso-3. Fly (Austin) 2012, 6, 80–92, doi:10.4161/fly.19695.

46. Obenchain, V.; Lawrence, M.; Carey, V.; Gogarten, S.; Shannon, P.; Morgan, M. VariantAnnotation: A Bioconductor Package for Exploration and Annotation of Genetic Variants. Bioinforma. Oxf. Engl. 2014, 30, 2076–2078, doi:10.1093/bioinformatics/btu168.

47. Fu, L.; Niu, B.; Zhu, Z.; Wu, S.; Li, W. CD-HIT: Accelerated for Clustering the next-Generation Sequencing Data. Bioinforma. Oxf. Engl. 2012, 28, 3150–3152, doi:10.1093/bioinformatics/bts565.

